# Accurate MS-based Rab10 phosphorylation stoichiometry determination as readout for LRRK2 activity in Parkinson’s disease

**DOI:** 10.1101/819607

**Authors:** Özge Karayel, Francesca Tonelli, Sebastian Virreira Winter, Phillip E. Geyer, Ying Fan, Esther M. Sammler, Dario R. Alessi, Martin Steger, Matthias Mann

## Abstract

Pathogenic mutations in the Leucine-rich repeat kinase 2 (LRRK2) are the predominant genetic cause of Parkinson’s disease (PD). They increase its activity, resulting in augmented Rab10-Thr73 phosphorylation and conversely, LRRK2 inhibition decreases pRab10 levels. However, there is no assay to quantify pRab10 levels for drug target engagement or patient stratification. We developed an ultra-sensitive targeted mass spectrometry (MS)-based assay for determining Rab10-Thr73 phosphorylation stoichiometry in human samples. It uses synthetic stable isotope-labeled (SIL) analogues for both phosphorylated and non-phosphorylated tryptic peptides surrounding Rab10-Thr73 to directly derive the percentage of Rab10 phosphorylation from attomole amounts of the endogenous phosphopeptide. We test the reproducibility of our assay by determining Rab10-Thr73 phosphorylation stoichiometry in human neutrophils before and after LRRK2 inhibition. Compared to healthy controls, neutrophils of LRRK2 G2019S and VPS35 D620N carriers robustly display 1.4-fold and 3.7-fold increased pRab10 levels, respectively. Our generic MS-based assay further establishes the relevance of pRab10 as a prognostic PD marker and is a powerful tool for determining LRRK2 inhibitor efficacy and for stratifying PD patients for LRRK2 inhibitor treatment.

## INTRODUCTION

Parkinson’s disease (PD) is the second most common neurodegenerative disease, and no disease-modifying therapies exist to date ^1,2^. Although most PD cases are idiopathic, mutations in several genes have been linked to familial forms of the disease ^3^. Among those, mutations in the Leucine-rich repeat kinase 2 (LRRK2) comprise the predominant genetic cause of PD and account for 1% of sporadic and 4% of familial cases worldwide, and much higher in some populations ^4^. At least six pathogenic missense mutations in LRRK2, including the most frequent G2019S substitution, have been identified ^5^ and several studies confirmed that these mutations increase its kinase activity ^5–8^. LRRK2-associated PD is clinically largely indistinguishable from idiopathic PD, suggesting that LRRK2 inhibition may be useful as disease-modifying therapy for a larger group of patients ^4^. Clinical trials with selective LRRK2 kinase inhibitors are ongoing and have already passed phase 1.

We have recently identified a subset of Rab GTPases (Rab3A/B/C/D, Rab5A/B/C, Rab8A/B, Rab10, Rab12, Rab29, Rab35, and Rab43) as LRRK2 substrates ^8,9^. Among these, Rab10 appears to be a key physiological kinase substrate ^8–14^. LRRK2 directly phosphorylates Rab10 at Thr73 and all previously tested pathogenic forms of LRRK2 enhance this phosphorylation ^8,9^. Intriguingly, the PD-associated D620N mutation of the retromer complex protein VPS35 also activates LRRK2 kinase activity, which in turn results in augmented Rab10 phosphorylation ^15^. Thus, multiple PD-associated factors are interconnected and dysregulation of a common LRRK2-Rab signaling pathway may be an underlying cause of PD.

The LRRK2 autophosphorylation site Ser1292 has been widely used for assessing LRRK2 kinase activity ^16–19^. However, its phosphorylation levels appear to be extremely low and there is no sensitive phospho-specific antibody available to reliably detect and quantify phosphorylation at this site ^6,20,21^. Instead, we recently developed several high-affinity antibodies for detecting pRab proteins in cells and in tissues ^14^. Among those, a highly specific clone detects pThr73 levels in human peripheral blood cells of (mutant) LRRK2 G2019S and VPS35-D620N carriers ^11,15^. While pRab10 levels were markedly increased in VPS35 D620N carriers, no statistically significant differences in pRab10 levels were detected by immunoblotting analysis when comparing controls to LRRK2-G2019S carriers in peripheral blood mononuclear cells (PBMCs) and neutrophils ^11,15^. The reproducible quantification of immunoblots, particularly the detection of small (<2) fold-changes is challenging. In contrast, mass spectrometry-based quantification has become the gold standard and has several advantages over traditional biochemical methods, as it is more specific and more precise. Importantly, it allows simultaneous detection of both the phosphorylated peptide and the total protein pool and hence calculation of the absolute fraction of the phosphorylated protein, also known as phosphorylation stoichiometry or occupancy ^22^.

There are several strategies differing in their accuracy, throughput, and applicability for measuring phosphorylation stoichiometry by MS ^22–27^. One way is to compare the intensities of modified and unmodified peptides by label-free proteomics ^27^. However, due to potential differences in ionization efficiencies, the MS signals of individual peptides cannot be directly compared with each other. Stoichiometry determination based on heavy-to-light ratios of stable isotope-labeled (SIL) analogues of phosphorylated and non-phosphorylated proteolytic peptides and their endogenous counterparts can in principle overcome this problem and result in a much more sensitive and precise readout for monitoring kinase activity ^22,23^. Alternatively, a SIL recombinant protein that is chemically or enzymatically phosphorylated can also be used as the spike-in standard ^28,29^. MS strategies determining stoichiometry that are combined with tailored targeted methods are especially suited for accurate quantification of low levels of a given phosphorylated analyte ^30^. In particular, instruments with an Orbitrap as a mass analyzer can be operated in targeted scan modes such as high resolution selected ion monitoring (SIM) or parallel reaction monitoring (PRM). In both methods, precursor ions are isolated with a narrow m/z range by a quadrupole mass filter before introduction into the Orbitrap analyzer, thereby providing increased signal to noise ratio for the ion of interest and allowing attomole-level limits of detection ^31,32^. As no fragmentation is involved in SIM, quantification of selected ions relies on the high resolution in the Orbitrap mass analyzer. In contrast, in PRM, fragment ions are used for quantification of the peptide. While this approach is more specific, the overall sensitivity of PRM can be lower, as the signal of a given precursor ion is distributed across multiple fragments, although the signal to noise ratio can be higher ^32^.

Here we describe an accurate and ultra-sensitive MS-assay for determining the Rab10-Thr73 phosphorylation stoichiometry in Parkinson’s disease. We evaluate our assay by measuring and comparing Rab10-Thr73 phosphorylation levels in healthy controls, idiopathic PD patients, and PD patients with defined genetic cause. Our assay enables the detection of subtle changes in Rab10 phosphorylation, which are beyond what is detectable by typical immunoassays merely targeting the phosphorylated species. We show that pRab10 stoichiometry when measured precisely using our assay can serve as a robust target engagement and patient stratification marker in clinical studies.

## RESULTS

### Rab10-pThr73 serves as a readout for LRRK2 activity in human peripheral blood neutrophils

To explore which Rab GTPases were expressed in human peripheral blood neutrophils and which of them could serve as a readout for LRRK2 activity in a quantitative MS-based assay, we first isolated neutrophils from whole blood using a negative selection approach. We assessed the recovery and the purity of the enriched cells by flow cytometry (Supplementary Fig. 1). Proteomic analysis resulted in 5,488 quantified proteins, for which we estimated the copy numbers per cell using the proteomic ruler approach ^33^ (Fig. 1A and Supplementary Table 1). LRRK2, the vesicular protein VPS35 and many of the Rab proteins that are LRRK2 substrates are found in the highest quartile of the abundance-ranked proteome (Fig. 1B). The bona-fide LRRK2 substrate Rab10 had the highest copy number among all detected Rab proteins with 1,820,000 copies per cell, suggesting that it could serve as an excellent marker for LRRK2 activity in PD (Supplementary Fig. 2).

**Fig. 1.**
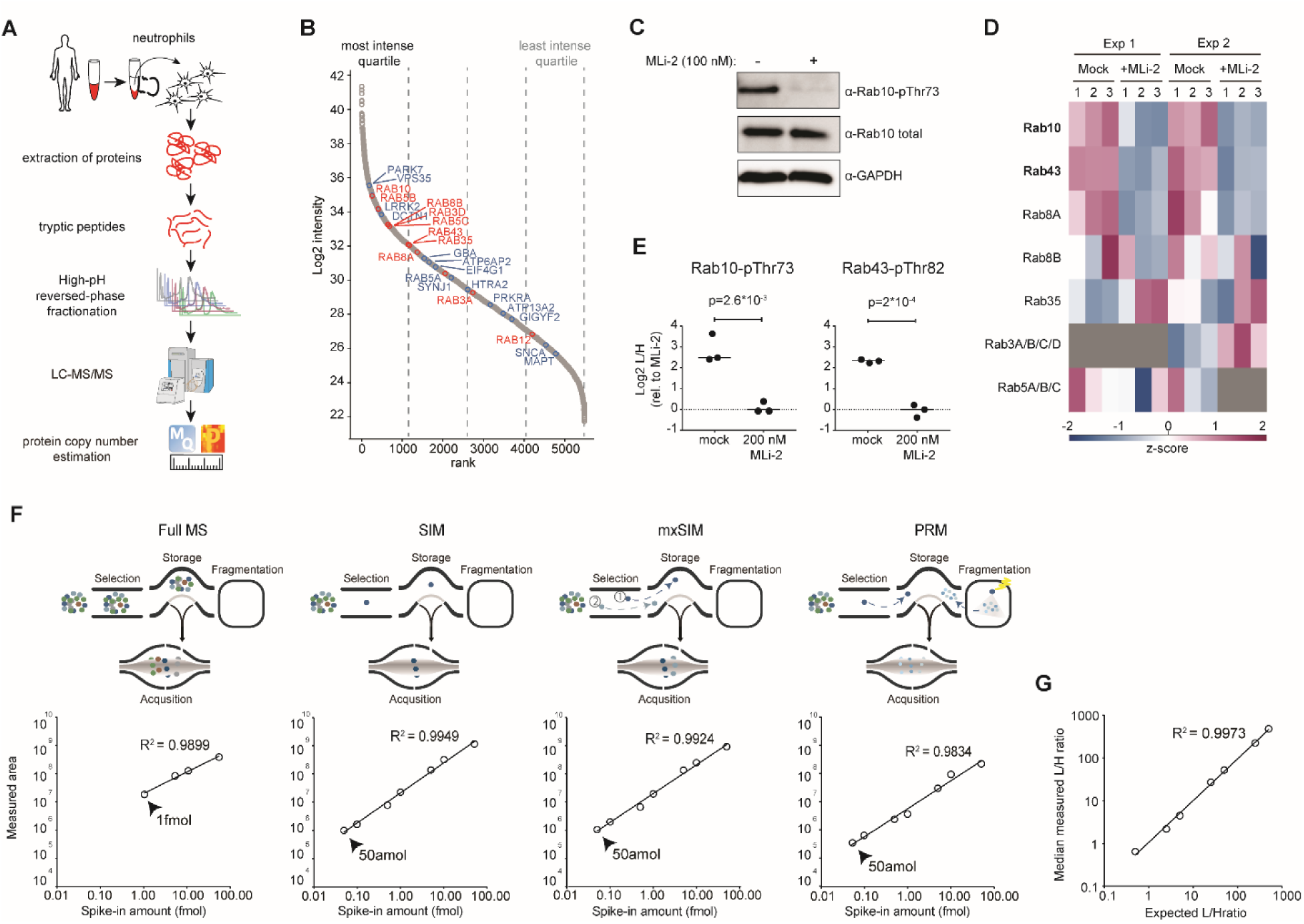
Rab10-pThr73 as a readout for LRRK2 activity in human peripheral blood neutrophils. (A) Workflow for global proteome analysis of human peripheral blood neutrophils. (B) Proteins ranked according to their abundances across the global neutrophil proteome. Quartiles are indicated with dashed lines. LRRK2 phosphorylated Rab proteins and other PD-associated proteins are highlighted in red and blue, respectively. (C) Western blot analysis of neutrophils (−/+200 nM MLi-2, 30 min) using anti-MJFF-pRAB10 (pThr73), Rab10 total and GAPDH antibodies. (D) Heat map of z-scored phosphopeptide intensities of LRRK2-phosphorylated Rab proteins from pRab immunoprecipitation of neutrophil lysates (−/+ 200 nM MLi-2, 30 min, 3 technical replicates, n =2). Values missing are shown in grey. (E) Targeted MS-quantified Rab10-pThr73 and Rab43-pThr82 peptide intensities in immunoprecipitations of individual Rab10 and Rab43 proteins from neutrophils (−/+ 200 nM MLi-2, 30 min). P-values were assessed using unpaired t-test analysis. (F) Detection limit (LOD) of SIL Rab10-pThr73 tryptic peptide (FHpTITTSYYR) with various acquisition methods; full MS, SIM, mxSIM and PRM. Linearity ranges of the dilution curves were obtained from singlets analysis of HeLa digest as background spiked with 10 amol to 50 fmol of the SIL Rab10-pThr73 phosphopeptide. (G) Limit of quantification (LOQ) for the Rab10-pThr73 tryptic peptide measured in mxSIM mode. 25 fmol of light phosphopeptide was mixed with variable amounts of its SIL counterpart (10 amol to 50 fmol) and spiked into a background of HeLa digest (3 technical replicates). Median ratios extracted from Skyline were plotted against the expected ratios.

To determine whether Rab10 is the only Rab family member that is phosphorylated by LRRK2 in neutrophils, we first treated freshly isolated cells with the selective LRRK2 inhibitor MLi-2 and confirmed the downregulation of Rab10-pThr73 by immunoblotting ^11^ (Fig. 1C). In the same lysates, we enriched phosphorylated Rab proteins with the previously described pThr-specific Rab antibody and subjected the eluates to LC-MS/MS analysis ^9^. We detected 19 Rab GTPases in total, of which 7 were previously shown to be LRRK2 targets ^9^ (Fig. 1D and Supplementary Fig. 3). However, in two independent pan-phospho Rab antibody pulldown experiments, only Rab10 and Rab43 phospho-protein levels decreased significantly upon MLi-2 treatment, indicating that these two Rabs are the endogenous targets in this system. To directly confirm LRRK2-mediated phosphorylation of Rab10 and Rab43 in neutrophils, we immunoprecipitated both Rab proteins and quantified Rab10-pThr73 and Rab43-pThr82 phosphopeptides by targeted mass spectrometry. Both phosphorylation sites were downregulated more than 4-fold after LRRK2 inhibitor treatment, demonstrating that the phosphorylation status of these Rabs can be used as readout for LRRK2 kinase activity (Fig. 1E).

The high abundance of Rab10 in neutrophils and the promise of pThr73 as a biomarker for Parkinson’s disease encouraged us to develop an ultra-sensitive targeted MS-based assay for quantifying the phosphorylated Rab10 peptide in human cells. To maximize the sensitivity, we explored a multiplexed SIM (mxSIM) setup on the latest generation linear quadrupole Orbitrap instrument (Q Exactive HF-X). In our method, the SIL analogue of the phosphorylated tryptic Rab10-Thr73 peptide acts as a sentinel peptide, as it can be spiked in in high amounts and elutes simultaneously with its endogenous light counterpart. Light (endogenous) as well as SIL counterpart phosphopeptides are consecutively isolated by narrow quadrupole isolation windows but simultaneously subjected to the Orbitrap mass analyzer. We set the maximum total ion accumulation time to 230 ms, but allocate 90% to the endogenous phosphopeptide, thus boosting its signal and increasing the sensitivity of our assay. To compare the sensitivity and the accuracy of (mx)SIM with regular full-MS scanning and with PRM, we mixed variable amounts of the SIL Rab10 phosphopeptide (10 amol-50 fmol) with 50 ng of a tryptic Hela digest and measured the SIL pRab10 peptide intensity using the different scan protocols. Relative to full-MS scanning, in which the entire mass range (300-1650 m/z) is analyzed, SIM, either multiplexed or not, provided a 20-fold increase in sensitivity with a limit of detection (LOD) of 50 amol (Fig. 1F). PRM performed equally well in terms of sensitivity, however, SIM had a higher quantification accuracy (R^2^ of 0.992 vs. 0.983). For this reason, and because it was sufficiently specific, we decided to develop a Rab10-pThr73 quantification assay based on mxSIM.

To determine the maximum heavy-to-light ratio and the limit of quantification (LOQ) of our method, we mixed 25 fmol of the light pRab10 peptide with variable amounts of its heavy counterpart (10 amol to 50 fmol) in 50 ng of HeLa digest. The correlation for mxSIM indicates an excellent reproducibility of quantification also in the multiplexed case (R^2^ = 0.997) (Fig. 1G). Due to the differential filling strategy, we accurately quantified heavy-to-light ratios of Rab10 phosphopeptides of up to 1:500 (25 fmol light and 50 amol heavy Rab10 peptide).

### mxSIM can precisely determine the Rab10-Thr73 phosphorylation stoichiometry

To evaluate the protein and peptide-centric approaches for determining Rab10-Thr73 phosphorylation stoichiometry, we expressed and purified Rab10 (1-175 aa) from an auxotroph *E.coli* strain, which allows for incorporation of SIL lysine and arginine into the newly synthesized protein ^29,34^. We phosphorylated 50% of the recombinant protein by LRRK2, as shown by intact mass analysis and bottom-up proteomics confirmed Thr73 as phosphorylation site (Supplementary Fig. 4A and 4B). This is a good proportion as both phosphorylated and non-phosphorylated peptides are needed as standards and we decided to use this SIL recombinant phosphoprotein for quantifying the percentage of pRab10 in cells. For this purpose, we immunoprecipitated HA-Rab10 from LRRK2-Y1699C-expressing HEK293 cells, either treated with MLi-2 or not, and mixed the enriched protein with our SIL standard before joint tryptic digestion (Supplementary Fig. 5A-B). We then derived the Rab10-Thr73 phosphorylation stoichiometry as described above, which turned out to be 66.62 ± 1.54%. Subsequently, we used the same measurement but calculated the occupancy with the peptide-centric approach, yielding 67.17 ± 1.10%. As both approaches gave nearly equal results, we decided to use the simpler peptide centric approach for all further calculations (Fig. 2B).

**Fig. 2.**
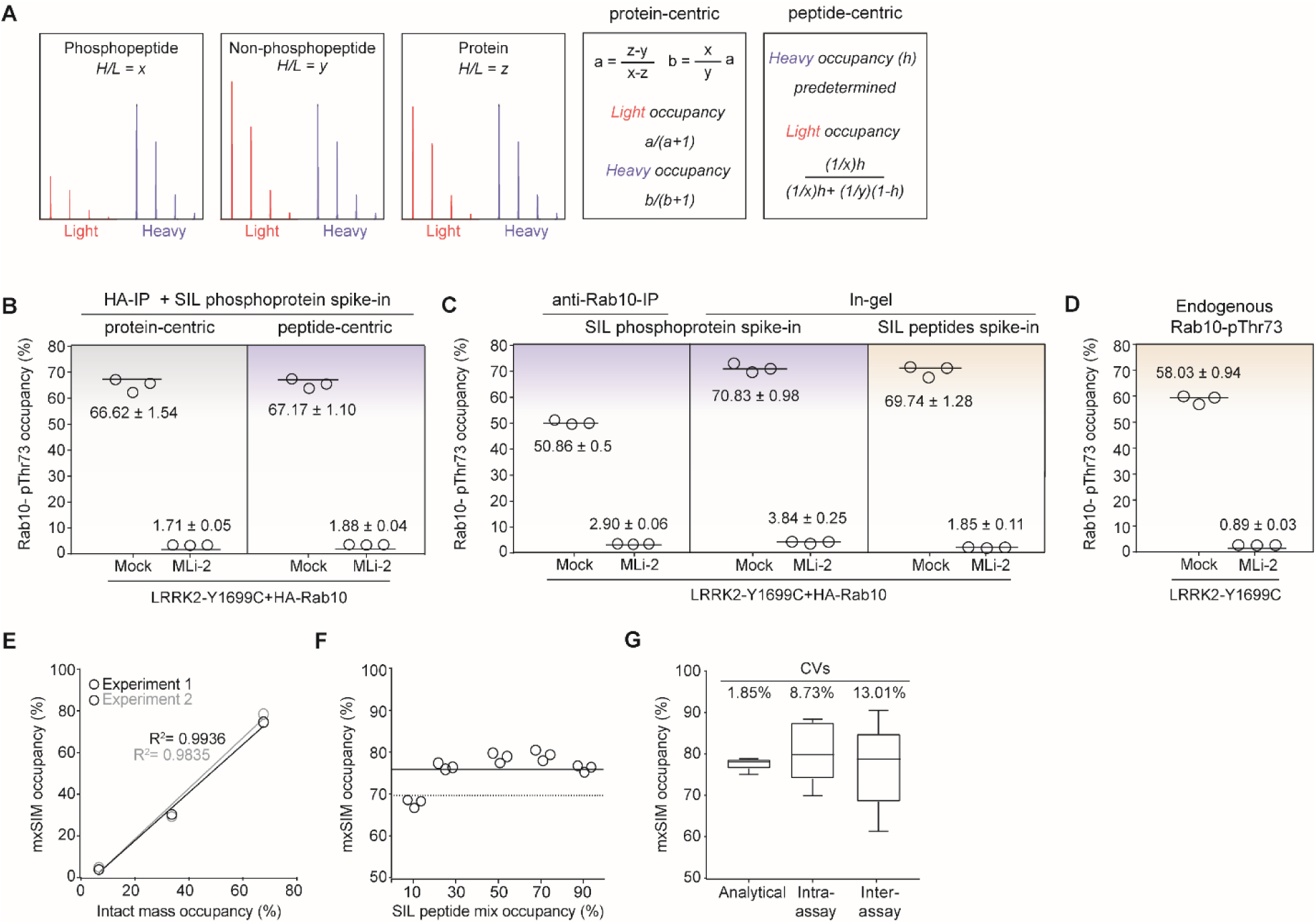
mxSIM precisely determines the Rab10-Thr73 phosphorylation stoichiometry. (A) Heavy-to-light ratios and the formulas used in protein- and peptide-centric approaches. (B) HA-Rab10-pThr73 occupancy in LRRK2-Y1699C-expressing HEK293 cells was determined using the protein- vs peptide-centric approaches after HA-IP. (C) HA-Rab10-pThr73 occupancy determined using the peptide-centric approach with either SIL phosphoprotein or SIL peptide standard spike-in after enrichment by anti-Rab10-IP or SDS-PAGE followed by in-gel digestion in HEK293 cells expressing LRRK2-Y1699C (−/+ 200 nM MLi-2, 60 min). (D) Endogenous Rab10-pThr73 occupancy determined using SIL peptide standards in mock and LRRK2-Y1699C expressing HEK293 cells (−/+ 200 nM MLi-2, 60 min). Samples from (B), (C) and (D) were analyzed in triplicates using the mxSIM method and the phosphorylation occupancies were presented as means ± SEM. (E) Benchmarking our method using Thr73-phosphorylated unlabeled recombinant Rab10 proteins (1-175 aa) as standards. Correlation of the median occupancies determined either by intact mass analysis or by the mxSIM method in triplicates for two independent experiments. (F) SIL phosphorylated and non-phosphorylated Rab10 peptides mixed in various ratios representing 10, 30, 50, 70 and 90% occupancies and spiked into standard phosphoprotein digest were measured using our mxSIM method in triplicates. The solid line represents the median occupancy (75.38 ± 1.46%) whereas the dashed line shows the estimated phosphorylation occupancy of the standard protein (70%) by intact mass analysis. (G) CVs were calculated by repeating MS measurements (analytical), and the workflow in the same gel (intra-assay) or in different gels (inter-assay) using the same phosphoprotein standard (n=6).

Detection of sub-stoichiometric, post-translationally modified peptides in complex mixtures by proteomics is challenging, even with very sensitive targeted methods, and requires one or more upfront enrichment steps. To select a suitable antibody for enriching Rab10, we compared an antibody recognizing the total protein with one recognizing the HA-epitope tag in HA-Rab10 expressing cells. Unexpectedly, the Rab10 directed antibody yielded significantly lower pThr73 occupancy as compared to the antibody directed against the epitope tag (50.86 ± 0.5% vs 67.17 ± 1.1%). A potential explanation for this discrepancy could be that the anti-Rab10 antibody preferentially recognizes the non-phosphorylated fraction of the total protein pool (Fig. 2B and C). To address this issue with a different approach that does not rely on antibodies, we separated the cell lysate mixed with the SIL phosphoprotein standard on SDS-PAGE, excised the region of ~15-30 kDa and digested the proteins using trypsin, followed by mxSIM analysis. This resulted in a measured Rab10-pThr73 occupancy of 70.83 ± 0.98%, which was strikingly similar to the occupancy obtained by the anti-HA immunoprecipitation approach (Fig. 2B-C). Finally, we used SIL phosphorylated and non-phosphorylated peptides for deriving the Rab10-Thr73 phosphosite occupancy and again found that 69.74 ± 1.28% of the protein was phosphorylated in the same cell lysate, an almost identical occupancy value as the one obtained through spike-in of the SIL phosphoprotein (Fig. 2C). We therefore decided to combine gel-separation, SIL peptide spike-in and the peptide-centric calculation approach. With these tools in hand, we extended our in-gel digestion workflow to endogenous Rab10-pThr73 occupancy determination. We found that 58.03 ± 0.94% of the protein was phosphorylated in LRRK2-Y1699C-transfected cells, whereas the treatment with MLi-2 almost completely abolished the phospho-occupancy (0.89 ± 0.03%) (Fig. 2D).

To further benchmark our assay, we incubated recombinant Rab10 (1-175 aa) with LRRK2 and stopped the phosphorylation reaction by adding the LRRK2 inhibitor HG-10-102-01 at defined time intervals. Next, we determined the percentages of the phosphorylated Rab10 proteins by intact mass spectrometry and compared these values to the occupancies obtained using our mxSIM method (Supplementary Fig. 6). This experiment revealed an excellent correlation between these methods in two independent experiments with R^2^ of 0.984 and 0.994 (Fig. 2E). Similarly, to specifically test the SIL peptide spike-in part of the approach, we mixed SIL phosphorylated and non-phosphorylated Rab10 peptides in 1:10, 1:3.3, 1:1, 3.3:1 and 10:1 ratios to mimic phosphosite occupancies of 10, 30, 50, 70 and 90%, respectively, and mixed them with our recombinant Rab10 phosphoprotein. We then derived the Rab10-pThr73 occupancy based on the heavy-to-light ratios of both phosphorylated and non-phosphorylated peptides (Fig. 2F). The calculated mean occupancy for our recombinant Rab10 phosphoprotein was 75.3 ± 1.46% and 70%, for mxSIM and intact mass analysis, respectively. This small difference is likely caused by the intensity measurements in the intact mass analysis (see Supplementary Material). Finally, we determined the analytical, the intra- and the inter-assay variabilities, which yielded excellent coefficients of variations (CVs) of 1.85%, 8.73% and 13.01%, respectively (Fig. 2G).

### In-gel digestion combined with mxSIM can detect subtle changes in Rab10-Thr73 phosphorylation within cells

Most pathogenic LRRK2 mutations, including R1441C/G/H, increase LRRK2 kinase activity and significantly stimulate Rab10 protein phosphorylation in mouse and human cells and tissues ^6–9^. To determine whether our assay was sufficiently accurate and robust for detecting small differences of LRRK2 activity in cells, we treated WT and LRRK2 R1441G knock-in mouse embryonic fibroblasts (MEFs) with increasing concentrations of MLi-2 and determined Rab10-Thr73 phosphorylation occupancies (Fig. 3A). In parallel, we controlled LRRK2 inhibitor efficacy by immunoblotting and probing for Rab10-pThr73 (Fig. 3B). Compared to WT, in which the Rab10-pThr73 occupancy was 12.12 ± 0.64%, we found a 2.45-fold increase in R1441G (29.66 ± 0.92%) (Fig. 3C). Rab10-pThr73 occupancy already decreased by 1.5-fold (18.75 ± 1.50%) upon treatment with 1 nM of MLi-2 and by almost 2-fold (15.38 ± 0.69%) with 3 nM of the inhibitor (Fig. 3B-C). The corresponding Rab10-pThr73 immunoblot signals also showed some decrease, however, reliable and precise quantification of these bands was difficult. To further extend our analysis, we determined a Rab10-pThr73 occupancy-based IC_50_ of 3 nM. This value is in in the 3-10 nM range estimated by our previous phos-tag Rab10 analysis (Fig. 3D) ^12^. Together, our results establish that our assay can reliably detect very small differences in phosphorylation occupancy in cultured cells.

**Fig. 3.**
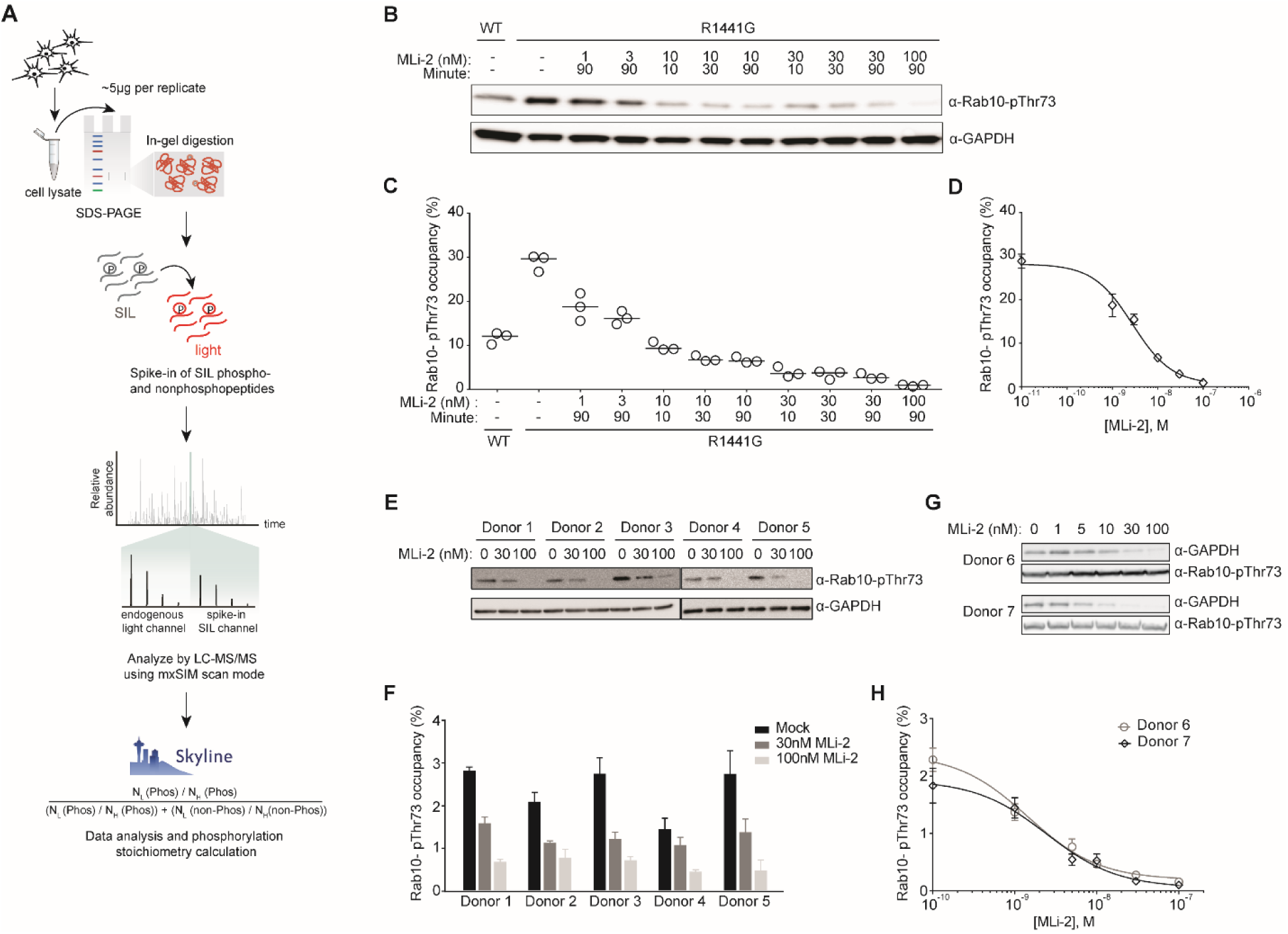
Our assay reliably determines the Rab10-pThr73 occupancy in cell lysates. (A) Workflow for the in vivo Rab10-pThr73 occupancy assay. (B) Immunoblotting of WT and R1441G knock-in MEFs treated with the indicated concentrations of MLi-2 for different intervals using monoclonal MJFF-pRAB10 (pThr73) and GAPDH antibodies. (C) Rab10-pThr73 occupancies were determined by mxSIM in the same lysates (n=3). (D) Dose–response curve of Rab10-pThr73 occupancy in R1441G knock-in MEFs to generate occupancy based-IC_50_ values for MLi-2. Each data point represents the median of triplicate measurements in the samples treated with MLi-2 for 90 min. Error bars represent SEM. (E) Immunoblotting of neutrophils isolated from healthy individuals (DMSO or 30 and 100 nM MLi-2 treated) using monoclonal MJFF-pRAB10 (pThr73) and GAPDH antibodies. (F) Rab10-pThr73 occupancies were determined by mxSIM in the same lysates. Error bars represent SEM. (G) Immunoblotting of the neutrophils isolated from healthy individuals and treated with the indicated concentrations of MLi-2 using monoclonal MJFF-pRAB10 (pThr73) and GAPDH antibodies. (H) Individual-specific dose–response curves of occupancies to generate occupancy based-IC_50_ for MLi-2.

Next, to determine whether our assay was sufficiently sensitive for quantifying the percentage of Rab10-pThr73 in human peripheral blood, we isolated neutrophils from five healthy volunteers. Upon treatment with either 30 nM or 100 nM of MLi-2, we found a decrease in Rab10-pThr73, as judged by immunoblotting (Fig. 3E). When we determined the phosphorylation stoichiometry by MS, we found very low levels of phosphorylated Rab10 protein in all individuals. In fact, the median occupancy was 2.34 ± 0.16% in DMSO-treated cells and this decreased to 1.25 ± 0.06% after 30 nM and to 0.6 ± 0.05% after 100 nM MLi-2 treatment (Fig. 3F). These occupancies correspond to 225,500 ± 15,500 (DMSO), 115,270 ± 6,190 (30 nM MLi-2) and 60,160 ± 4,600 (100 nM MLi-2) phosphorylated Rab10 molecules per cells (Supplementary Table 2).

Finally, we performed a MLi-2 dose-response experiment (1-100 nM) using neutrophils from two donors and determined their occupancy based-IC_50_ values. These were very similar in both donors (2.44 and 3 nM), further demonstrating the high reproducibility of our method even *in vivo* (Fig.3G and H).

### PD patients show an increased Rab10-Thr73 phosphorylation stoichiometry

The frequency of the common LRRK2 G2019S mutation is 1% of patients with sporadic PD and 4% of patients with hereditary PD and even higher in certain populations such as Ashkenazi Jews and North African Berbers ^4^. To investigate the central question of whether our assay can stratify PD patients with elevated LRRK2 activity in clinical trials, we analyzed Rab10-pThr73 levels in neutrophils of four LRRK2-G2019S carriers, together with an equal number of healthy controls in a blinded experimental setup (see Methods). Each neutrophil population was treated with DMSO or 100 nM MLi-2 and all samples were subjected to quantitative immunoblot analysis for Rab10-pThr73 levels, revealing a clear reduction in Rab10 phosphorylation upon LRRK2 inhibition. Total Rab10 levels did not differ between treated and untreated, and healthy controls and heterozygous G2019S mutation carriers (Fig. 4A). The median occupancy of Rab10-pThr73 were 3.06 ± 0.14% in healthy controls whereas they increased by about 1.4-fold to 4.29 ± 0.59% in G2019S carriers. Although our data shows that Rab10-pThr73 is increased in G2019S carriers, the observed differences did not reach statistical significance due to the large intergroup variability and the small cohort size (Fig. 4C). In MLi-2 treated samples, occupancy decreased 5-fold on average for both groups (Fig. 4B).

**Fig. 4.**
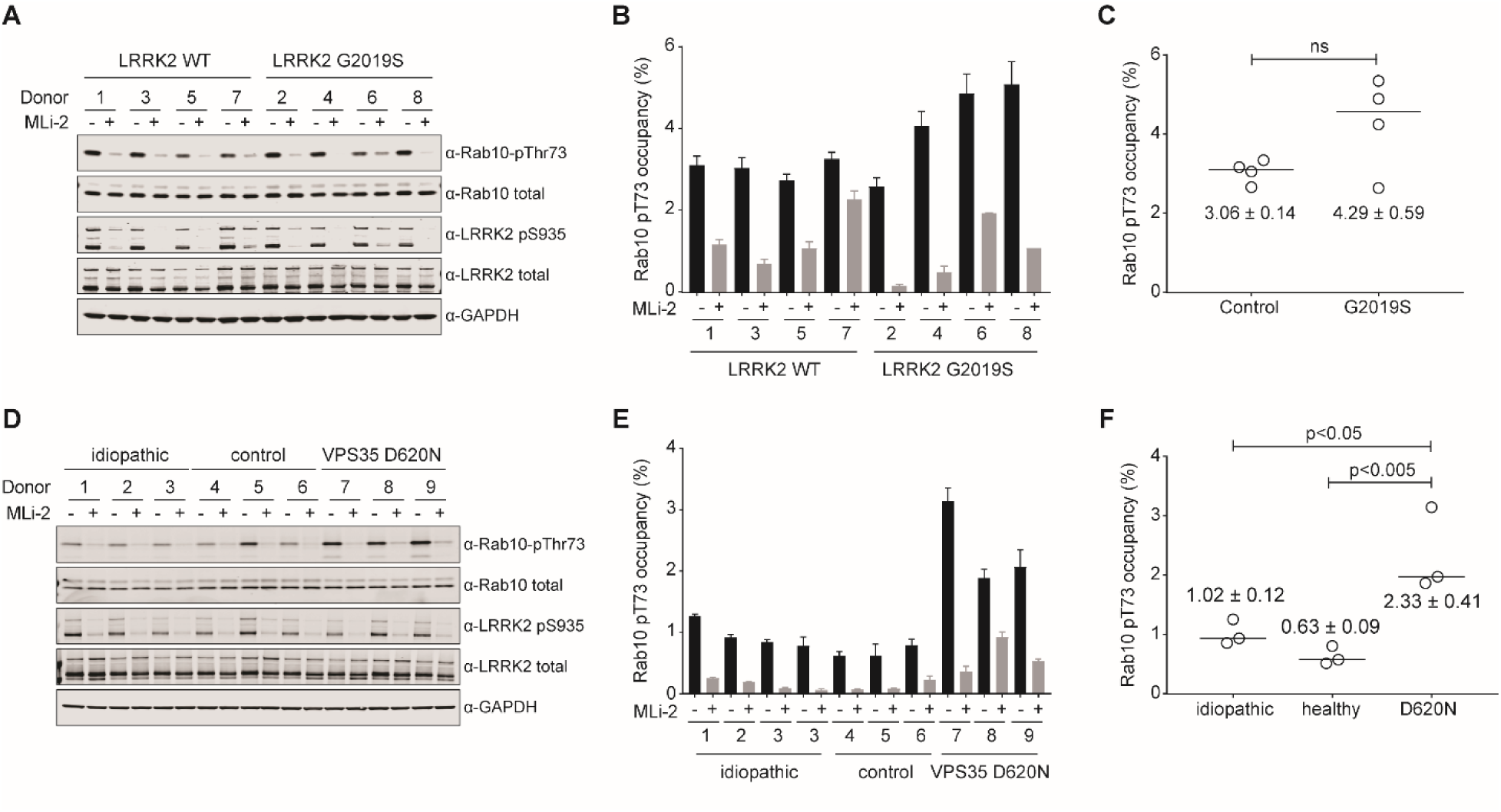
Our assay reveals changes in the Rab10-pThr73 occupancy in PD patient-derived neutrophils. (A) Immunoblotting of neutrophils isolated from four PD patients with the G2019S LRRK2 mutation and four healthy controls (−/+100 nM MLi-2) using anti total LRRK2, pSer935 LRRK2, total Rab10, MJFF-pRAB10 (pThr73) and GAPDH antibodies. (B) Blind application of our mxSIM assay to the same lysates (n=3). Error bars represent SEM. (C) Quantification of Rab10-pThr73 occupancy between controls and PD patients with the G2019S LRRK2 mutation. Each data point represents the median of triplicate measurements in untreated samples. One-way ANOVA with Bonferroni’s multiple comparisons test was applied and the occupancies are presented as means ± SEM; ns: non-significant, *P < 0.05, **P < 0.005. (D) Immunoblotting of neutrophils isolated from three controls, three idiopathic PD patients and three PD patients with a heterogeneous VPS35 D620N mutation (−/+ 200 nM MLi-2) using anti total LRRK2, pSer935 LRRK2, total Rab10, MJFF-pRAB10 (pThr73) and GAPDH antibodies. (E) Blind application of our assay in the same lysates (n=3). Error bars represent SEM. (F) Quantification of Rab10-pThr73 occupancy between controls, idiopathic cases and PD patients with heterogeneous VPS35 D620N. The significant analysis was done as explained earlier in (C).

Our previous work demonstrated that VPS35 controls LRRK2 activity and that the D620N substitution resulted in increased LRRK2 activity as assessed by monitoring phospho-Rabs levels ^15^. To test if our assay can determine LRRK2 activity in PD patients with the VPS35 D620N mutation, we measured Rab10-Thr73 phosphorylation occupancies in neutrophils isolated from three heterozygous patients, as well as three age-matched idiopathic PD patients and three non-PD controls. We treated the neutrophils from each subject with either DMSO or 200 nM MLi-2 before cell lysis. In accordance with our previous work, immunoblot analysis revealed elevated Rab10-pThr73 levels in VPS35 D620N patients (Fig. 4D). LRRK2-pSer935 levels were also monitored by immunoblot analysis in the lysates to confirm the efficacy of MLi-2 treatment. The median occupancies were 0.63 ± 0.09%, 1.02 ± 0.12% and 2.33 ± 0.41% in healthy controls, idiopathic and heterozygous VPS35 D620N patients, respectively, and decreased 6.9-fold on average in MLi-2 treated cells (Fig. 4E). While the 1.6-fold increase of Rab10-pThr73 in idiopathic PD was not statistically significant, the absolute Rab10-pThr73 occupancy in heterozygous VPS35 D620N patients was significantly higher compared to controls and idiopathic cases (Fig. 4F). Together, these experiments demonstrate that our assay can reveal patients with higher LRRK2 kinase activity and suggest that it can be applied to stratify patients.

## DISCUSSION

Here, we have established an accurate and ultra-sensitive MS-based assay for determining the percentage of Rab10-Thr73 phosphorylation in samples collected from PD patients. We confirmed Rab10-Thr73 phosphorylation as a surrogate parameter for LRRK2 activity and specifically applied our assay to monitor LRRK2 activity in neutrophils. These cells contain relatively high levels of both LRRK2 and Rab10 and their isolation is straightforward, although robust isolation with minimal protease activation is important ^11^. As a next step, it will be interesting to investigate whether pRab10 can be detected in bodily fluids such as cerebrospinal fluid (CSF) and urine. Large collections of these bio-fluids that are readily accessible could be used to further establish pRab10 as a bona-fide biomarker for PD.

We find that the levels of Rab10-Thr73 phosphorylation are stable but very low in human cells (<5%). This is also true for LRRK2-G2019S mutation carriers where activation of the LRRK2 kinase is modest while individuals with a PD-associated VPS35 D620N mutation demonstrate much higher Rab10-Thr73 phosphorylation reflecting a much greater catalytic effect on the LRRK2 kinase. This indicates that changes in pRab10 levels are critical for PD pathogenesis and the very low percentage of pRab10 may explain the age-dependent, reduced penetrance of the LRRK2 G2019S mutation. It will therefore be interesting to compare pRab10 percentages of matched disease-manifesting and non-manifesting LRRK2 G2019S mutation carriers at younger and older ages.

Our results show that minimal changes in the total pool of a phosphorylated protein within the cells can trigger pathogenesis. As a next step, stratification for Rab10-pThr73 levels in PD could help identify individuals with increased LRRK2 kinase activity who would most likely benefit from LRRK2 kinase inhibitor treatment. However, our mass spectrometric assay is completely generic and not restricted to a particular disease. In the future, it could be applied to study any other kinase-substrate relation in clinical reasearch. For example, phosphosite occupancies of prominent oncogenic factors could give important information on how phosphorylation stoichiometry influences tumorigenesis.

Our assay utilizes separation of Rab10 by SDS-PAGE, which is followed by in-gel digestion. This strategy enriches Rab10 significantly and allows quantifying Rab phosphorylation with high sensitivity. In the application to very large cohorts, a limitation of our assay is its throughput. For example, the ability to immunoprecipitated total Rab10 with specific antibodies that recognize both GDP and GTP bound Rab10 with equal affinities would be highly desirable and considerable effort is currently being invested in producing these tools^14^. Alternatively, non-hydrolysable GTP analogues could be used to enrich GTP Rab proteins before MS analysis.

We determined pRab10 occupancy in neutrophils of 14 healthy controls, three idiopathic PD patients, four LRRK2 G2019S and three VPS35 D620N mutation carriers. As expected, compared to the matched idiopathic PD patients and healthy controls, we found a higher fraction of phosphorylated Rab10 in both LRRK2 G2019S and VPS35 D620N, although in the mutant LRRK2 group the difference did not reach statistical significance. This is readily explained by the small number of patients, the large intragroup variability and the very moderate fold-changes induced by LRRK2 G2019S. We expect that analysis of larger cohorts and inclusion of patient samples with higher LRRK2 activity, such as the R1441G/C mutations, will further establish pRab10 occupancy as a bona-fide marker of PD. It will be exciting to investigate if these patients with increased pRab10 levels also benefit from LRRK2 inhibitor treatment.

In summary, our generic and very sensitive MS-based assay accurately measures phosphorylation stoichiometry of any given substrate. It can be used for ongoing clinical LRRK2 inhibitor studies, in which the target engagement, dosing and efficacy of the compounds needs to be evaluated. We envision that our assay can also be applied to further investigate the role of LRRK2 in the PD, potentially revealing new upstream players of LRRK2 pathway, and determining whether they have a potential in treating PD.

## MATERIAL AND METHODS

### Reagents

MLi-2 (cat# 5756) was purchased from Tocris Bioscience. DIFP, HA-agarose and trypsin were from Sigma and LysC was from Wako. Microcystin-LR was from Enzo Life Sciences. Complete protease and phosphatase inhibitor tablets were from Roche. SIL peptides in absolute quantities (AQUA) were purchased from Sigma-Aldrich.

### Antibodies

Anti-GAPDH (#5174), anti-HA (#3724), and anti-Rab10 (#8127) were from Cell Signaling Technologies. Rabbit monoclonal antibody for total LRRK2 (UDD3, ab133518), Rab10 (T73) antibody [MJF-R21] and polyclonal phospho-Rab (initially raised against Rab8) antibody [MJF-R20] that efficiently immunoprecipitated multiple LRRK2 phosphorylated Rab proteins were purchased from Abcam (Burlingame, California). They were custom-made by Abcam in collaboration with the Michael J. Fox Foundation (Burlingame, California) ^14,35^.

### Plasmids

HA-Rab10 (DU44250) and Flag-LRRK2-Y1699C (DU13165). Full datasheets and reagents are available on https://mrcppureagents.dundee.ac.uk/.

### Stable isotope labeling and purification of the Rab10 protein standard

Buffer A: 50 mM HEPES pH 8.0, 500 mM LiCl, 1 mM MgCl2, 100 μM GDP, 1 mM TCEP and Elution buffer: Buffer A + 500 mM imidazole

To obtain unlabeled Rab10 standard, human His-tagged Rab10 (residues 1-175) construct from Rai et al.^36^ was expressed in *E. coli* BL21 (DE3) harboring the GroEL/S plasmid and protein expression was induced with isopropyl β-D-1-thiogalactopyranoside overnight. Cells were lysed by sonication in a buffer containing 50 mM HEPES pH 8.0, 500 mM LiCl, 1 mM MgCl2, 100 μM GDP, 1 mM TCEP (Buffer A) supplemented with several protease inhibitors including 30 μM Antipain and 50 μM Chymostain. Proteins were purified by Ni-NTA affinity chromatography. Briefly, bound proteins eluted with an imidazole gradient (25mM-500mM) and imidazole was removed by washing with 500 μM ATP containing Buffer A. Further purification was done by ion-exchange chromatography (Q-Sepharose) followed by size exclusion chromatography using a Superdex 200 column. Peak fractions containing recombinant protein were pooled. Identity and purity of the standard protein were assessed by Maldi-TOF MS and SDS-PAGE. To obtain labeled Rab10 standard, an auxotrophic expression strain for arginine and lysine was used ^29,34^. Cultures were grown in PA5052 minimal autoinduction media containing heavy Arg10 and Lys8 and cells were harvested for purification of SIL Rab10 protein standard as described earlier.

### *In vitro* LRRK2 kinase assays

Recombinant LRRK2 G2019S (Invitrogen, PV4881) and Rab proteins were incubated at 30°C for 30 min in kinase assay buffer (100 mM Tris– HCl, pH 7.5, 50 mM MgCl2, 5 mM EGTA, 1 mM GDP, 10 mM DTT, 25 mM β-glycerol phosphate, 5mM Sodium orthovanadate, and 50 μM ATP). The reaction was terminated by addition of 2 μM HG-10-102-0.

### Cell culture and transfection

HEK293 and MEFs (WT and LRRK2-R1441G) cells were cultured in Dulbecco’s modified Eagle medium (Glutamax, Gibco) supplemented with 10% fetal calf serum, 100 U/ml penicillin and 100 µg/ml streptomycin. Transient transfections were performed 48 hours prior to cell lysis using polyethylenimine PEI (Polysciences). Transfected cells were subjected to DMSO or MLi-2 (dissolved in DMSO) treatments at concentrations and periods of time as indicated in each figure legend. All cells were tested for mycoplasma contamination and overexpressing lines were verified by western blot analysis.

### Study participants and blood sample collection

For setting up and validating our assay, we recruited volunteers from within the Department of Proteomics and Signal Transduction at the Max Planck Institute of Biochemistry who kindly donated blood for our study. The data shown in Figures 1 and 3 and Supplementary Figures 1–3 are derived blood samples from healthy donors, which provided a written informed consent, with prior approval of the ethics committee of the Max Planck Society.

### Neutrophil isolation, characterization, treatments and lysis

Neutrophils were isolated directly from freshly drawn EDTA whole blood by immune-magnetic negative isolation using the MACSxpress® Neutrophil Isolation Kit (Miltenyi Biotec, Cat# 130-104-434). For 8 ml of blood, one vial of ‘Isolation Cocktail’ (magnetic beads) from the neutrophil isolation kit, delivered as a lyophilized pellet, was reconstituted by adding 0.25 mL of Buffer A and 0.25 mL of Buffer B in order. Then, the cocktail was mixed by gently pipetting and added to 8 ml of whole blood in 15 ml falcon tube. The blood sample containing the cocktail was gently mixed by inversion and incubated at room temperature for 5 min. The falcon tube was next placed into the magnetic field of the MACSxpress Separator (# 130-098-308) for 15 min. The magnetically labeled cells (non-neutrophils) adhere to the wall of the tube while the aggregated erythrocytes sediment to the bottom. The supernatant (~ 7 ml) containing the enriched neutrophils was carefully pipetted into a new 15 ml falcon tube, avoiding touching the magnetic beads attached to the sides of the falcon tube as well as the red blood cells at the bottom of the tube. The supernatant containing the isolated neutrophils was centrifuged at 300×g for 10 min at room temperature (acceleration and deceleration is both 5 using a Beckman Coulter Allegra X-15R Centrifuge). To ensure the removal of erythrocytes, the pelleted neutrophil cells were resuspended in 10 ml of 1× Red Blood Cell Lysis Solution (# 130-094-183), incubated for 10 min at room temperature and centrifuged at 300×g for 10 min at room temperature. To assess the purity and recovery of the enriched neutrophils we fluorescently stained cells with CD14-PerCP, CD15-PE, CD16-APC, and CD193-FITC by flow cytometry. Cell debris, dead cells were excluded from the analysis based on scatter signals and DAPI. Finally, the cell pellet was resuspended with 10 ml room temperature RPMI 1640 media by gentle pipetting. At this stage, purified cells were subjected to MLi-2 (dissolved in DMSO) treatment at concentrations and periods of time as indicated in each figure legend. An equivalent volume of DMSO was added to negative control samples. Following treatment, cells were pelleted through centrifugation at 500 g for 5 min. Cells were then resuspended in 5 ml of ice-cold PBS and centrifuged again at 500 g for 5 min. The supernatant was carefully removed by pipetting. Cells were lysed in 300 μl of ice-cold NP-40 buffer (50 mM Tris–HCl, pH 7.5, 1% (v/v) Triton X-100, 1 mM EGTA, 1 mM sodium orthovanadate, 50 mM NaF, 0.1% (v/v) 2-mercaptoethanol, 10 mM 2-glycerophosphate, 5 mM sodium pyrophosphate, 0.1 µg/ml microcystin-LR, 270 mM sucrose, 0.5 mM DIFP (Sigma, Cat# D0879) supplemented with protease and phosphatase inhibitors (Roche)). Lysate was clarified by centrifugation at 16,000 rpm for 15 min at 4°C after a liquid nitrogen freeze-thaw cycle. Protein concentration was measured using Bradford assay (Thermo Scientific), snap-frozen and stored at −80°C.

For the experiments shown in Figure 4, neutrophil lysates derived from either idiopathic PD patients, PD patients carrying a heterozygous LRRK2 G2019S or VPS35 D620N mutation or non-PD controls were used that had been previously used for publication ([11] and [15] respectively). All procedures were performed in compliance with the local ethics review boards and all participants provided informed consent. These lysates were subjected to MS analysis in a blinded experimental set-up, with the identity of the lysates only being revealed after completion of the MS analysis.

### Immunoblot analysis

Cell lysates were mixed with 4× SDS–PAGE loading buffer [250 mM Tris–HCl, pH 6.8, 8% (w/v) SDS, 40% (v/v) glycerol, 0.02% (w/v) Bromophenol Blue and 4% (v/v) 2-mercaptoethanol] to final total protein concentration of 1 µg/µl and heated at 85°C for 10 min. Samples were loaded onto NuPAGE Bis-Tris 4–12% gel (Life Technologies) and electrophoresed at 180 V for 1 h with MOPS SDS running buffer followed by transfer onto the nitrocellulose membrane (GE Healthcare, Amersham Protran Supported 0.45 µm NC) at 100 V for 90 min on ice in the transfer buffer (48 mM Tris–HCl and 39 mM glycine). Membrane was then cropped into pieces: from top of the membrane to 75 kDa to incubate with rabbit anti-LRRK2 UDD3 antibody, from 75 to 30 kDa to incubate with rabbit anti-GAPDH antibody and from 30 kDa to the bottom of the membrane to incubate with rabbit monoclonal antibodies for anti-Rab10-pThr73 or anti-Rab10. Antibodies diluted in 5% (w/v) BSA in TBS-T to a final concentration of 1 µg/ml. All blots were incubated in primary antibody overnight at 4°C. Prior to secondary antibody incubation, membranes were washed three times with TBS-T [20 mM Tris/HCl, pH 7.5, 150 mM NaCl and 0.2% (v/v) Tween 20] for 10 min each. Membranes were incubated with secondary antibody multiplexed with goat anti-rabbit diluted in 5% (w/v) non-fat dry milk (NFDM) in TBS-T (1:5000 dilution) for 1 h at room temperature. They were next washed with TBS-T three times for 10 min. Protein bands were detected using an ECL solution (Amersham ECL Western Blotting Detection Reagents (GE Healthcare)) and the ImageQuant LAS 4000 imaging system. For the immunoblots shown in Figure 4A and 4D, the membranes were developed using the LI-COR Odyssey CLx Western Blot imaging system and signal quantified using the Image Studio software.

### Cells lysis and pull-downs

Cells were lysed in either NP-40 buffer (50 mM Tris-HCl, pH 7.5, 120 mM NaCl, 1 mM EDTA, 6 mM EGTA, 20mM NaF, 15 mM sodium pyrophosphate and 1 % NP-40 supplemented with protease and phosphatase inhibitors (Roche)) or Triton X-100 buffer (50 mM Tris–HCl, pH 7.5, 1% (v/v) Triton X-100, 1 mM EGTA, 1 mM sodium orthovanadate, 50 mM NaF, 0.1% (v/v) 2-mercaptoethanol, 10 mM 2-glycerophosphate, 5 mM sodium pyrophosphate, 0.1 µg/ml microcystin-LR, 270 mM sucrose, 0.5 mM DIFP (Sigma, Cat# D0879) supplemented with protease and phosphatase inhibitors (Roche)). Lysates were clarified by centrifugation at 16,000 rpm for 15 min at 4°C after a liquid nitrogen freeze-thaw cycle. Protein concentrations were measured using Bradford assay (Thermo Scientific), snap-frozen and stored at −80°C. For HA pulldowns, lysates were incubated with HA-agarose resin for 2 h (25 µl resin per 100 ug of lysates). For immunoprecipitation using total Rab10 or phospho-Rab, lysates were incubated with antibodies in manufacturer’s recommend dilutions or concentrations overnight at 4°C and subsequently incubated with 25 μl of Protein-A/G-agarose beads for 2 h at 4°C. Unspecific binders were washed twice with matching lysis buffer and twice with 50 mM Tris-HCl (pH 7.5). SIL phosphorylated Rab10 protein was mixed with the beads at this step if used as standrad. Washes were followed by on-bead digestion overnight at 37°C with trypsin (~500 ng/pulldown in urea buffer [2M urea dissolved in 50 mM ammonium bicarbonate] or SDC buffer [1% (w/v) SDC in 100 mM Tris-HCL pH 8.5]). The resulting peptides were processed as described in ‘LC-MS/MS sample preparation’.

### In-Gel digestion protocol

Cell lysates were mixed with 4 × SDS/PAGE sample buffer (250 mM Tris/HCl, pH 6.8, 8% (w/v) SDS, 40% (v/v) glycerol, 0.02% (w/v) Bromophenol Blue and 4% (v/v) 2-mercaptoethanol) and heated at 85°C for 5 min. SIL Rab10 phosphoprotein was mixed with samples at this step if used as standard. 20-30 μg sample (5 μg per MS analysis) was loaded onto NuPAGE Bis-Tris 4–12% gel (Life Technologies) and electrophoresed at 180 V. After SDS-PAGE, gel was washed once with deionized water and stained with 0.1% Coomassie Blue R250 in 10% acetic acid, 40% methanol and 60% deionized water for 20 min and subsequently destained by soaking for at least 2 h in 10% acetic acid, 40% methanol, and 60% deionized water with at least two changes of the solvent (until the background is nearly clear). Gel band corresponding to 20-30 kDa is excised and chopped into smaller pieces (~ 1 × 1 mm) and placed in clean 1.5 ml tubes. Gel pieces are washed two or three times with 50% 50 mM ABC / 50% EtOH for 20 min at RT and then completely dehydrated by incubating for 10 min in absolute EtOH. Samples were dried in a speed-vac for 10 min (45°C) until the gel pieces were bouncing in the tube. Gel pieces were rehydrated in 300 µl of 1% (w/v) SDC buffer (10 mM TCEP, 40 mM CAA, trypsin in 100 mM Tris-HCL pH 8.5) per sample and placed at 37°C overnight. The next day, 300 µl of isopropanol buffer (1% TFA in isopropanol) was added and shaken for 10 min and spin down. The liquid was transferred into a fresh tube. Further 200 µl of isopropanol buffer was added to the gel pieces and shaken vigorously for 20 min at RT. It was combined with that from the previous step. Samples were directly loaded onto SDB-RPS stage tips and processed as described in ‘LC-MS/MS sample preparation’.

### Human Neutrophil Proteome Digestion, In-StageTip Purification and Fractionation

The neutrophil cell pellet was prepared with the iST Kit for proteomic sample preparation (P.O. 00001, PreOmics GmbH). In brief, this involved denaturation, alkylation, digestion and peptide purification. Reduction of disulfide bridges, cysteine alkylation and protein denaturation was performed at 95°C for 10 min. After a 5 min cooling step at room temperature, trypsin and LysC were added to the mixture at a ratio of 1:100 micrograms of enzyme to micrograms of protein. Digestion was performed at 37°C for 1 h. An amount of 20 μg of peptides was loaded on two 14-gauge SDB-RPS StageTip plugs. Samples were directly loaded onto SDB-RPS stage tips processed as described in ‘LC-MS/MS sample preparation’. Clean peptides were fractionated using the high-pH reversed-phase ‘Spider fractionator’ into 24 fractions as described previously to generate deep proteomes ^37^.

### LC-MS/MS sample preparation

StageTips ^38^ were prepared by inserting two 16-gauge layers of a SDB-RPS matrix (Empore) into a 200 μl pipette tip using an in-house prepared syringe device as described previously ^39^. The StageTips were centrifuged using an in-house 3D-printed StageTip centrifugal device at 1500 g. The acidified peptides (1% TFA v/v) were loaded onto the StageTips that were later washed with 1% TFA in isopropanol and subsequently 2% ACN/0.2% TFA. Elution was performed using 60 μl of 50% ACN/1.25% NH_4_OH or 80% ACN/1.25 % NH_4_OH. Eluates were collected in PCR tubes and dried using a SpeedVac centrifuge (Eppendorf, Concentrator plus) at 60°C. Peptides were resuspended in buffer A* (2% ACN/0.1% TFA) and briefly sonicated (Branson Ultrasonics) before LC/MS-MS analysis. To calculate absolute Rab10-pThr73 occupancy, we spiked SIL phosphorylated and non-phosphorylated counterpart peptides into samples at this step.

### LC-MS/MS measurements

Peptides were loaded on a 20 or 50 cm reversed phase column (75 µm inner diameter, packed in house with ReproSil-Pur C18-AQ 1.9 µm resin (Dr. Maisch GmbH)). Column temperature was maintained at 60°C using a homemade column oven. An EASY-nLC 1200 system (Thermo Fisher Scientific) was directly coupled online with the mass spectrometer (Q Exactive HF-X, Thermo Fisher Scientific) via a nano-electrospray source, and peptides were separated with a binary buffer system of buffer A (0.1% formic acid (FA)) and buffer B (80% acetonitrile plus 0.1% FA), at a flow rate of 300 nl/min. Peptides were eluted with a 45 min gradient of 5-60% buffer B (0.1% (v/v) FA, 80% (v/v) ACN). After each gradient, the column was washed with 95% buffer B for 5 min.

The mass spectrometer was programmed to acquire in targeted scan mode in which every full scan with resolution 60,000 at 200 m/z (3 × 10^6^ ions accumulated with a maximum injection time of 20 ms) was followed by two multiplexed selected ion monitoring (mxSIM) scans employing multiplexing degree of two to record both light (endogenous) and heavy counterpart simultaneously for either phosphorylated or non-phosphorylated Rab10-pT73 tryptic peptides. Each SIM scan covered a range of m/z 150–2000 with resolution 120,000 (10^5^ ions accumulated with a maximum injection time of 230 ms for both light and heavy counterparts, 1.4 m/z isolation window and 0.4 m/z isolation offset). m/z values of doubly-charged Rab10-pT73 tryptic (FHpTITTSYYR) light and heavy peptides (Arginine labeled (13C, 15N)) were defined as follows: 684.8028 and 689.8070 whereas doubly-charged Rab10 non-phosphorylated tryptic (FHTITTSYYR) light and heavy peptides (Arginine labeled (13C, 15N)) were targeted at m/z of 644.8197 and 649.8238.

For the LOD experiment (Fig. 1), the mass spectrometer was programmed to acquire in either full scan mode alone or SIM or PRM combined with a full scan. Full scan acquisition was performed with a resolution of 120,000 at 200 m/z (3 × 10^6^ ions accumulated with a maximum injection time of 230 ms) to cover the scan range of 350-1,650 m/z. SIM acquisition was performed using a resolution of 120,000 at 200 m/z, isolation windows of 1.4 m/z with 0.4 m/z offset, target AGC values of 2 × 10^5^, and a maximum injection time of 230 ms. PRM acquisition was performed using a resolution of 60,000 at 200 m/z,, isolation windows of 1.4 m/z with 0.4 m/z offset, target AGC values of 2 × 10^5^, and a maximum injection time of 130 ms. Fragmentation was performed with a normalized collision energy of 27.

The neutrophil proteomes were analyzed using an LC-MS instrumentation consisting of an EASY-nLC 1200 system (Thermo Fisher Scientific) combined with a Q Exactive HF Orbitrap (Thermo Fisher Scientific) and a nano-electrospray ion source (Thermo Fisher Scientific). The purified peptides were separated on a 50 cm HPLC column (75 µm inner diameter, in-house packed into the tip with ReproSil-Pur C18-AQ 1.9 µm resin (Dr. Maisch GmbH)). Of each of the 24 fractions around 0.5 µg peptides were analyzed with a 45 min gradient. Peptides were loaded in buffer A (0.1% FA, 5% DMSO (v/v)) and eluted with a linear 35 min gradient of 3-30% of buffer B (0.1% FA, 5% DMSO, 80% (v/v) ACN), followed by a 7 min increase to 75% of buffer B and a 1 min increase to 98% of buffer B, and a 2 min wash of 98% buffer B at a flow rate of 450 nl/min. Column temperature was kept at 60°C by a Peltier element containing in-house developed oven. MS data were acquired with a Top15 data-dependent MS/MS scan method (topN method). Target values for the full scan MS spectra was 3 × 10^6^ charges in the 300-1,650 m/z range with a maximum injection time of 55 ms and a resolution of 120,000 at m/z 200. Fragmentation of precursor ions was performed by higher-energy C-trap dissociation (HCD) with a normalized collision energy of 27. MS/MS scans were performed at a resolution of 15,000 at m/z 200 with an ion target value of 5 × 10^4^ and a maximum injection time of 25 ms.

For intact mass analysis, proteins were loaded on a reversed-phase column (Phenomenex AerisTM 3,6 µm Widepore C4 100 mm × 2.1 mm inner diameter, 200 Å pore size). An Agilent 1100 HPLC system was coupled online with the mass spectrometer (microTOF, Bruker Daltonik) and masses were recorded from 800-3,000 m/z. Proteins were separated with a binary buffer system of buffer A (0.05% TFA in H_2_O, pH 2.0) and buffer B (0.05% TFA in can, pH 2.0) at a flow rate of 250 nl/min. Proteins were eluted with a gradient of 20-80% buffer B in 20 min. After each gradient, the column was washed with 95% buffer B for one minute and 20% buffer B for 3 min. Data were processed using the CompassTM ‘DataAnalysis’ software from Bruker Daltonik, deconvoluted with ‘MaximumEntropy’ and an instrument resolving power of 10,000.

### Data analysis and phosphorylation stoichiometry calculations

For the calculation of absolute Rab10-pThr73 occupancies, raw MS data were processed using Skyline ^40^ which is an open source software project and can be freely installed. The details and tutorials can be viewed on the website (http://proteome.gs.washington.edu/software/skyline). Raw files were directly imported into Skyline in their native file format. After data import, graphical displays of chromatographic traces for the top two isotopic peaks were manually inspected for proper peak picking of MS1 filtered peptides. All quantitation performed for phosphorylation occupancy calculations in this study were done on the precursor ion level. Only the most abundant first two peaks of the isotope cluster were used for quantitation. Peptide areas for the non-phosphorylated tryptic Rab10 peptide (FHTITTSYYR, m/z 644.8197++) and the phosphorylated tryptic Rab10 peptide (FHpTITTSYYR, m/z 684.8028++) with their R=13C615N4 heavy analogues were extracted to derive light-to-heavy ratios. The absolute quantification was determined by comparing the abundance of the known SIL internal standard peptides with the native peptides. The phosphorylation stoichiometry/occupancy is calculated by taking the ratio of the total amount of phosphorylated fraction to the total amount of both phosphorylated and non-phosphorylated forms, which is always represented as percentage (%). All details for occupancy calculations in this study are provided in the Supplementary Table 2.

For the deep proteome of human neutrophils, raw MS data were processed using MaxQuant version 1.5.6.8 ^40,41^ with an FDR < 0.01 at the peptide and protein level against the Human UniProt FASTA database (2017). Enzyme specificity was set to trypsin, and the search included cysteine carbamidomethylation as a fixed modification and N-acetylation of protein and oxidation of methionine as variable modifications. Up to two missed cleavages were allowed for protease digestion, and peptides had to be fully tryptic.

Bioinformatic analyses in this study were performed with Perseus (www.perseus-framework.org) ^42^, Microsoft Excel and data visualized using GraphPad Prism (GraphPad Software) or RStudio (https://www.rstudio.com/). Proteomics raw data have been deposited to the ProteomeXchange Consortium (http://proteomecentral.proteomexchange.org) via the PRIDE partner repository with the data set identifier PXD015219.

## ACKNOWLEDGEMENT

We thank Sabine Suppmann, Leopold Urich, Stephan Uebel, Stefan Pettera, Martin Spitaler, Nagarjuna Nagaraj, Victoria Sanchez and Antonio Piras from the MPIB Biochemistry Core Facility. We also thank Florian Meier, Susanne Kroiss, Gabriele Sowa, Igor Paron, Christian Deiml, Johannes B. Mueller and all the members of the department of Proteomics and Signal Transduction for their assistances and helpful discussions. We value the contributions of Shalini Padmanabhan, Marco Baptista (all from the Michael J. Fox Foundation for Parkinson’s research), Kalpana Merchant (TransThera Consulting) and Suzanne Pfeffer (University of Stanford). We are grateful that the patients and healthy volunteers kindly donated blood for the present study. We would like to thank Thomas Gasser (University of Tubingen, Tubingen, Germany) and Alexander Zimprich (Medical University of Vienna, Wien, Austria) for providing access to LRRK2 [G2019S] and VPS35 [D620N] mutation carriers respectively, as well as controls and idiopathic PD patients, that donated blood for neutrophil isolation. GroEL/S and C terminal truncated construct of Rab10 (1-175) were kindly provided by Dr. Matthias Müller from the Max Planck Institute of Molecular Physiology in Dortmund. LRRK2 R1441G MEFs were kindly provided by Dr Shu-Leong Ho (Division of Neurology, Department of Medicine, University of Hong Kong).

---

**Supplementary Fig 1.**
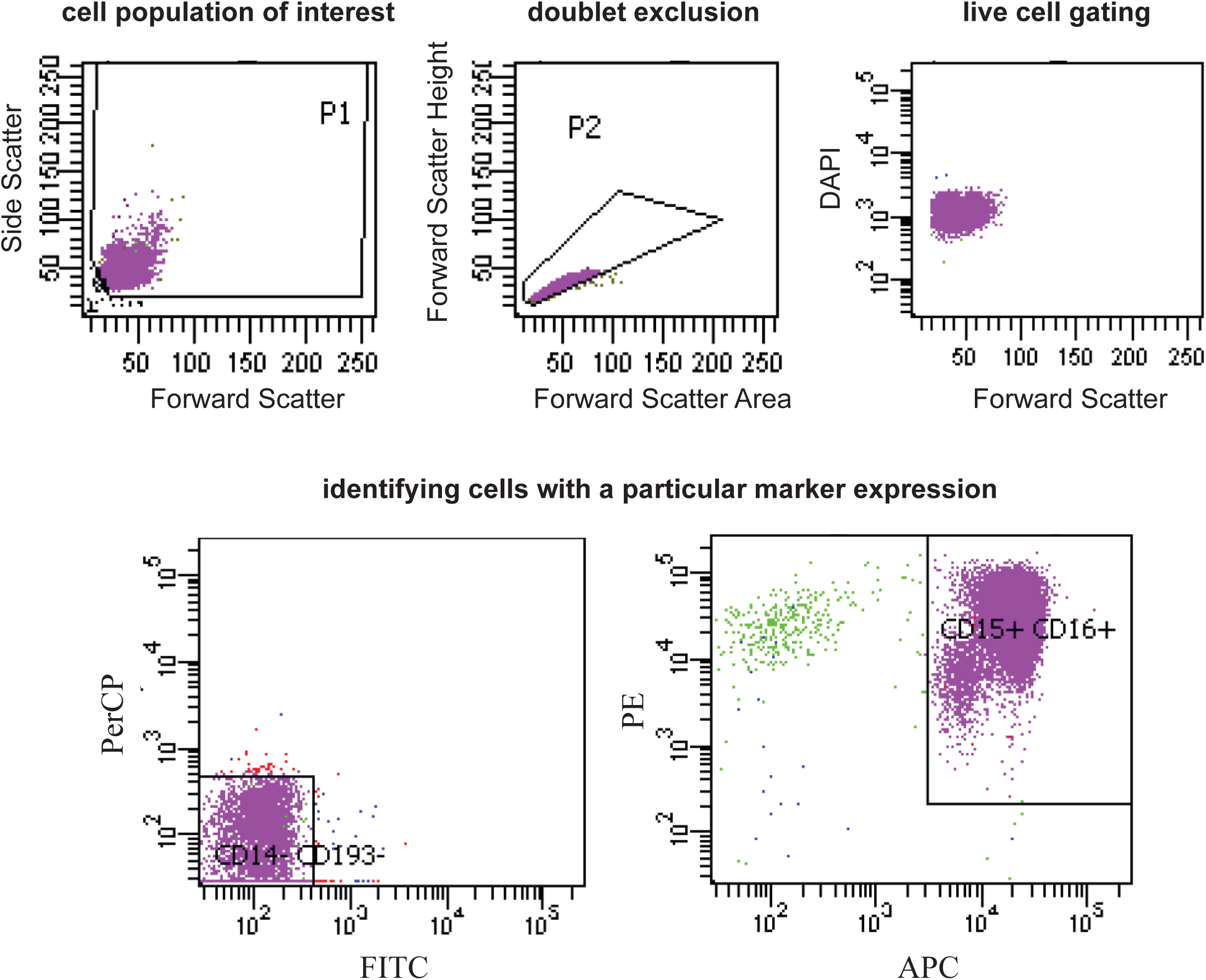
Flow cytometry analysis determining the purity of the isolation method and viability of isolated neutrophils. Plots and population gating shown is for the donor used in the experiment in Figure 1C-D. Cell debris, non-leukocytes, and dead cells were excluded from the analysis based on scatter signals and DAPI. Purity was assessed by staining with CD14-PerCP (monocytes and macrophages), CD15-PE (granulocytes, monocytes, neutrophils and eosinophils), CD16-APC (neutrophils, macrophages and eosinophils), and CD193-FITC (eosinophils). The population which is CD15+ and CD16+ but CD14- and CD193-represents the neutrophils. The isolated cells had a viability >96% and a purity>98%.

**Supplementary Fig. 2.**
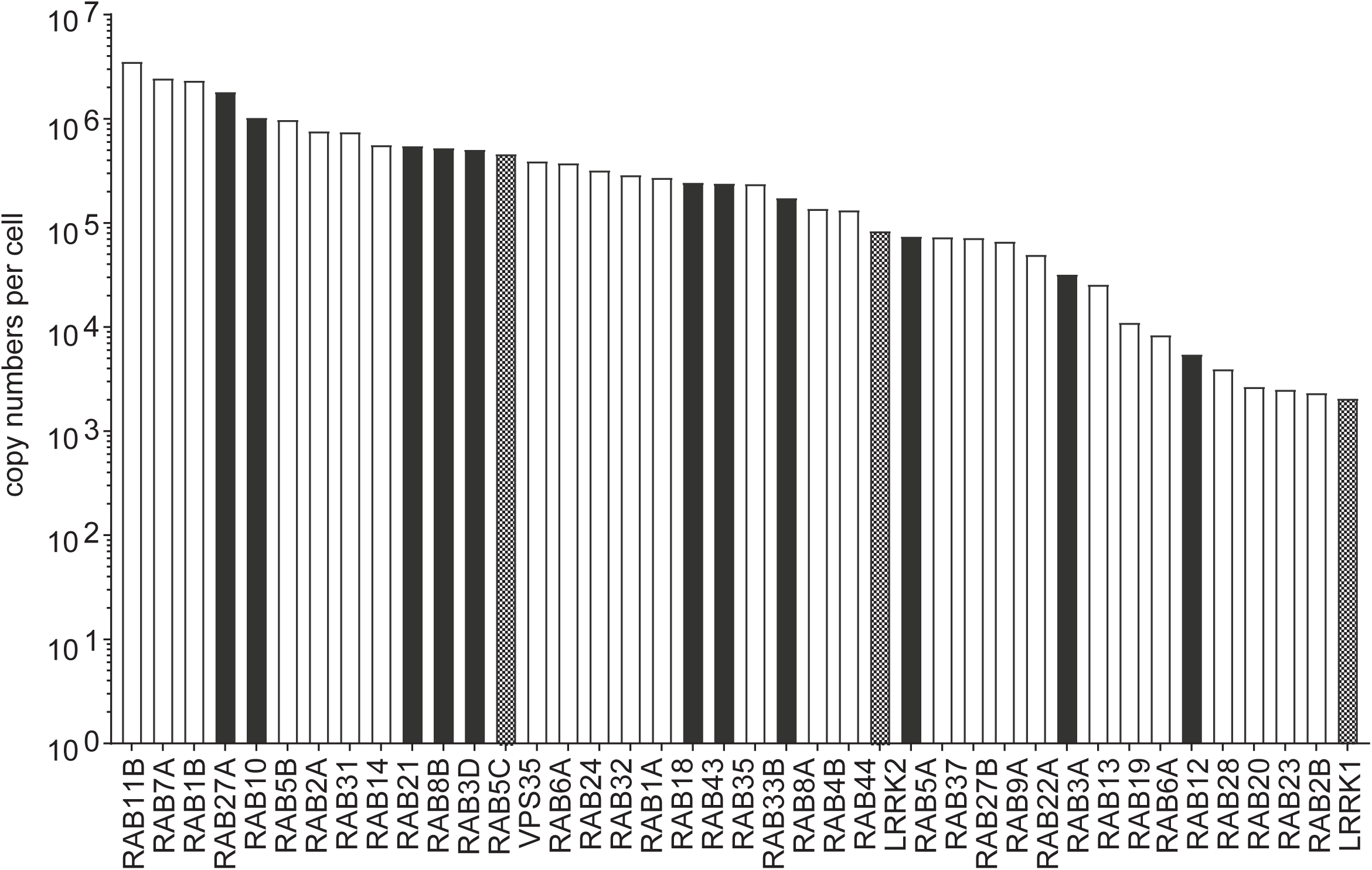
The copy numbers of the protein products of PD-associated genes, LRRK2 and VPS35, and Rab GTPases in a human neutrophil based on the cumulative histone amount considered proportional to the expected DNA amount per cell were estimated using the proteomic ruler approach (Wiśniewski et al., 2014), implemented in Perseus software (Tyanova et al., 2016). Black bars show the Rab proteins which were shown to be LRRK2 targets where as grey bars show LRRK1 and LRRK2 proteins.

**Supplementary Fig. 3.**
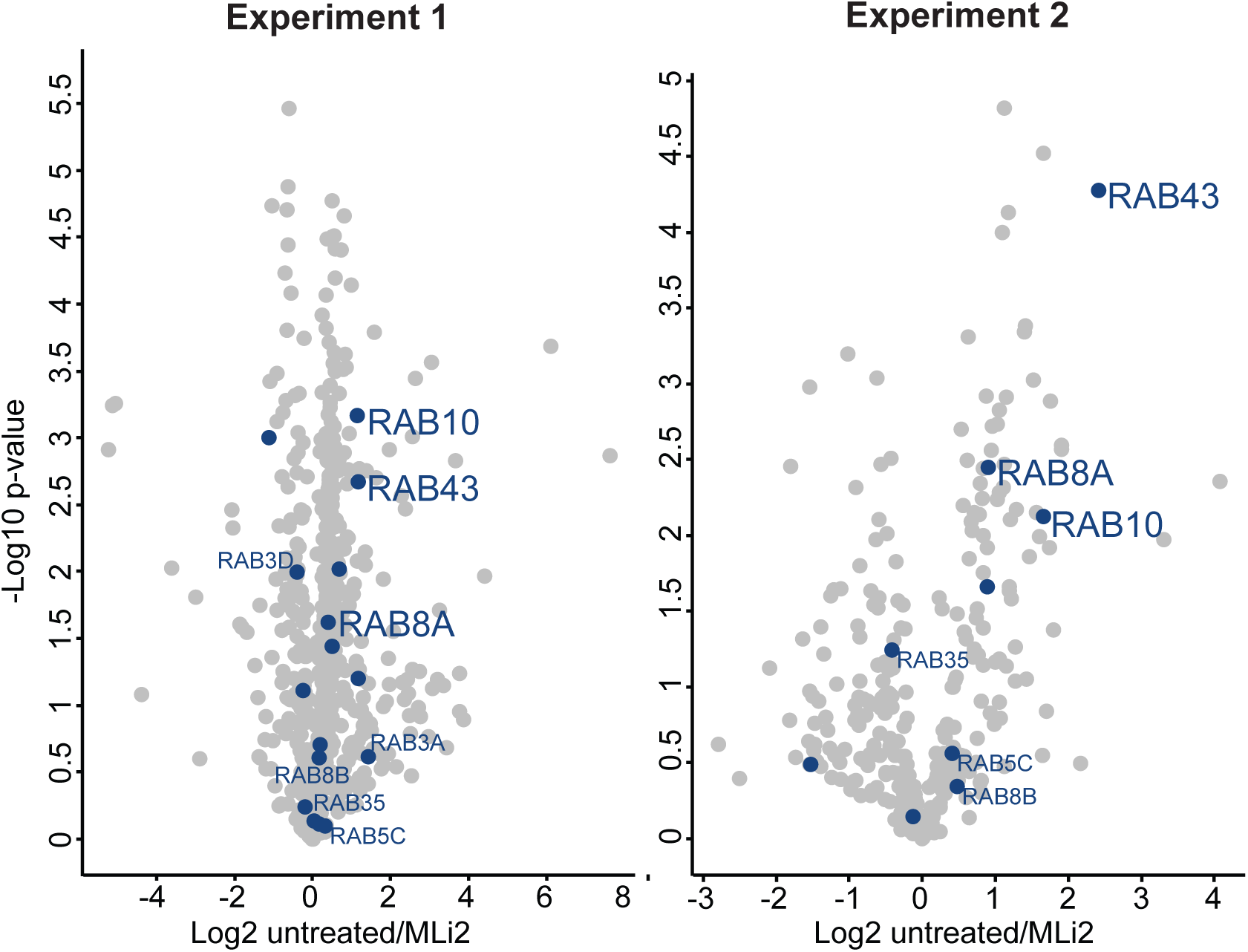
LRRK2 dependent Rab GTPase phosphorylations in human neutrophils. Volcano plots of the pRab immunoprecipitations (untreated:right and MLi-2-treated:left). All Rab proteins identified were highligted in blue and LRRK2 substrates were also labeled.

**Supplementary Fig. 4.**
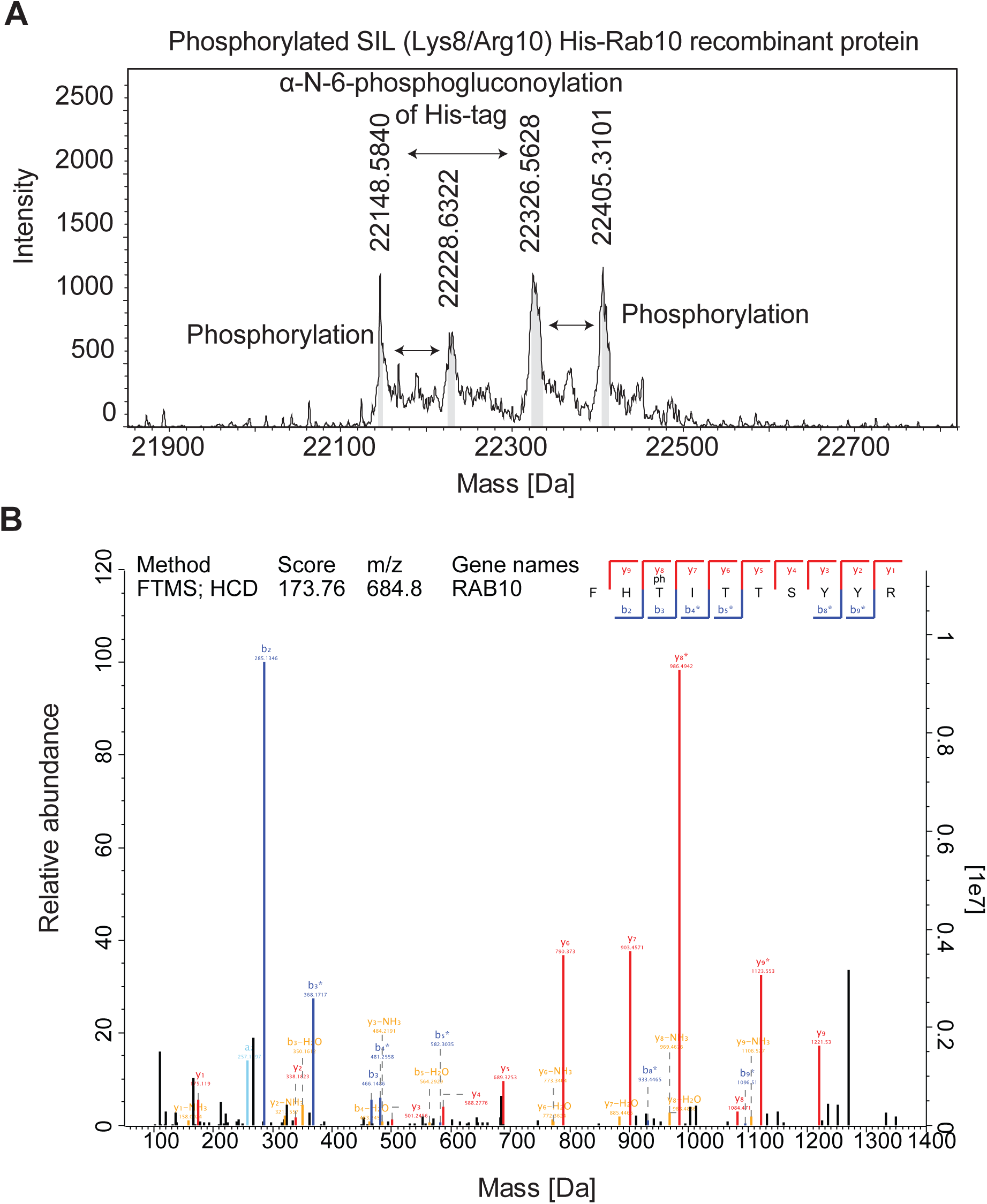
Phosphorylated SIL (Lys8/Arg10) His-Rab10 recombinant protein measured by both intact mass analysis and bottom-up proteomics. His-tagged Rab10 protein expressed and stable isotopically labeled (Lys8/Arg10) in E.coli was phosphorylated by the truncated (950-2527) recombinant human LRRK2-G2019S (Thermo Fisher, PV4881) in vitro. (A) The intact mass of the protein was analyzed by mTOF. We observed the mass of the protein (residues 1-175, m=22,148 Da) with a second peak with a mass difference of 178Da (m=22326 Da), indicative of α-N-6-phos-phogluconoylation of His-tag (Geoghegan, K. F. et al., 1999) and 80Da mass increases in both peaks due to phosphorylation. (B) LC-MS/MS analysis after digestion confirmed phosphorylation at T73. The collision-induced dissociation (CID) fragmentation spectrum with the Andromeda score (Cox et al., 2011) for the tryptic pT73-Rab10 peptide are shown.

**Supplementary Fig. 5.**
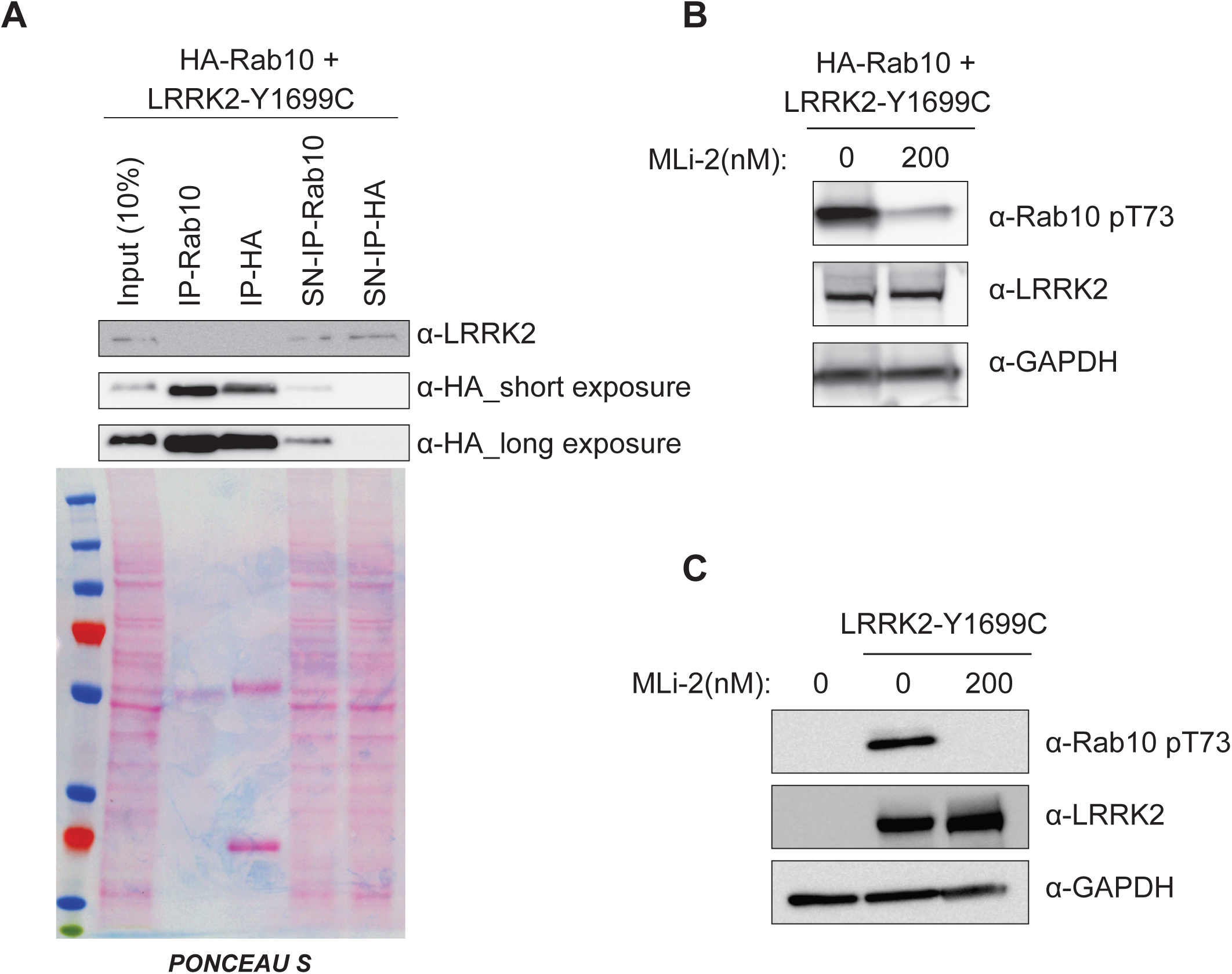
Rab10 protein either ectopically expressed or endogenously present in HEK293 cells with LRRK2-Y1699C expression. (A) Immunoblotting of HA-tagged Rab10 protein immunoprecipitation using HA agarose beads or anti-Rab10 antibody in HEK293 cells expressing LRRK2-Y1699C. The heavy chains can be seen in Ponceau staining. Input: whole cell lysate, IP: eluate, SN: supernatant. Immunoblottings of (B) HA-Rab10 and LRRK2-Y1699C and (C) mock and LRRK2-Y1699C expressing HEK293 cells (−/+ 200 nM MLi-2, 60 min) using total LRRK2, MJFF-pRAB10 (pThr73) and a loading control GAPDH antibody.

**Supplementary Fig. 6.**
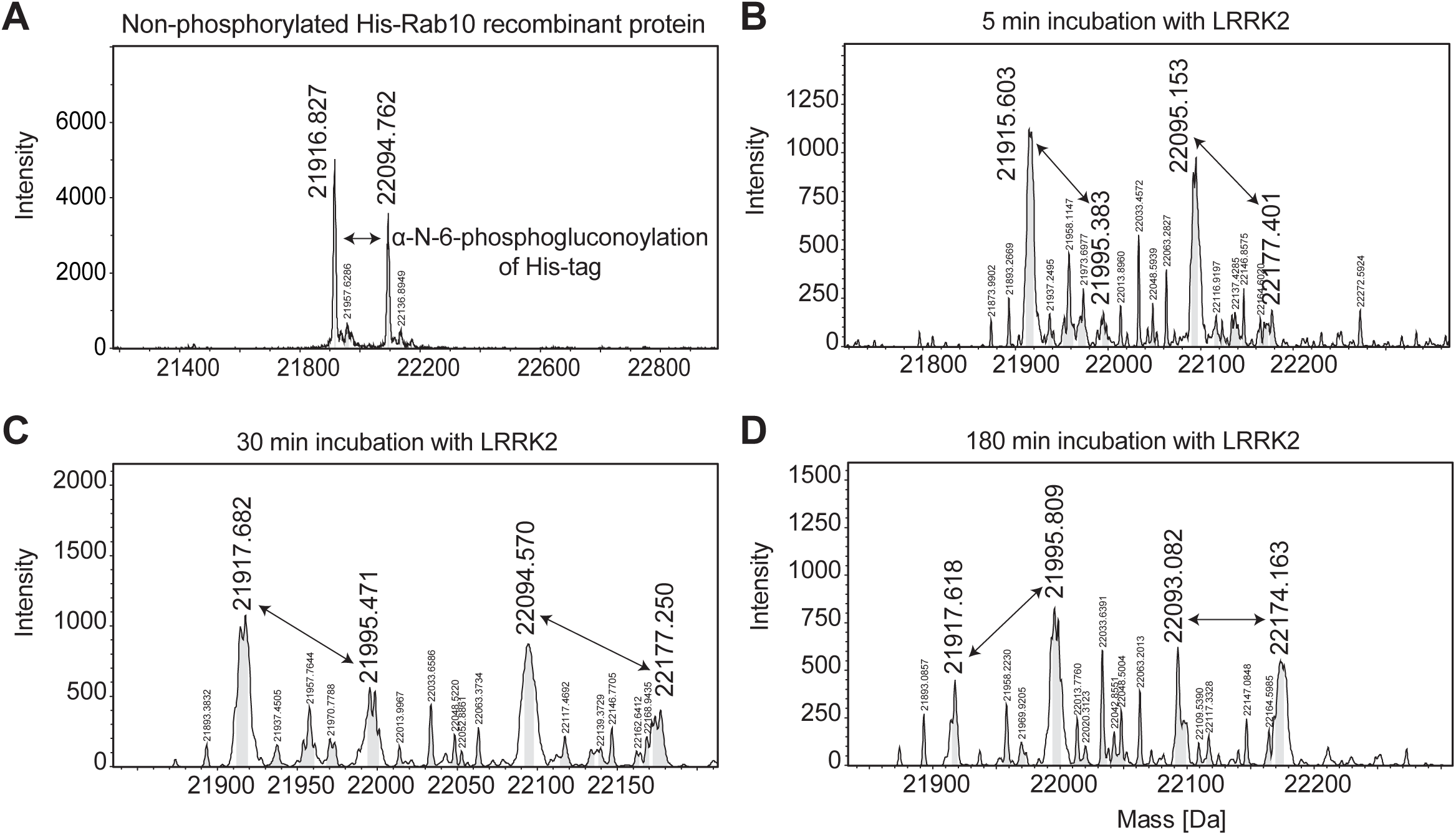
Intact mass analysis of non-phopshorylated and phosphorylated unlabeled His-Rab10 recombinant protein. (A) Rab10 protein expressed in E.coli was analyzed by mTOF. We observed the actual mass of the protein (m=21,916 Da) and α-N-6-phosphogluconoylated version (m=22,094 Da) (Geoghegan, K. F. et al., 1999). It was phosphorylated by the truncated (950-2527) recombinant human LRRK2-G2019S (Thermo Fisher, PV4881) in vitro. The reaction was stopped by the addition of 2 μM of HG-10-102-0, a selective LRRK2 inhibitor, after (B) 5 min, (C) 30 min and (D) 180 min to obtain phosphoprotein standards with different occupancies. Intact mass analysis by mTOF confirmed the phosphorylation events by identification of 80Da mass shifts for both unmodified and α-N-6-phosphogluconoylated peaks.

